# Porous yet Dense matrices: using ice to shape collagen 3D cell culture systems with increased physiological relevance

**DOI:** 10.1101/2022.03.13.484060

**Authors:** Cleo Parisi, Bénédicte Thiébot, Gervaise Mosser, Léa Trichet, Philippe Manivet, Francisco M. Fernandes

**Affiliations:** Sorbonne Université, UMR 7574, Laboratoire de Chimie de la Matière Condensée de Paris, 75005, Paris, France; Biobank Lariboisière/Saint Louis – Centre de Ressources Biologiques, Lariboisière Hospital, 75010, Paris, France; CY Cergy Paris Université, Université d’Evry, Université Paris-Saclay, CNRS, LAMBE, F-95000, Cergy, France; INSERM UMR1141 NeuroDiderot, Université de Paris, 75019, Paris, France

## Abstract

Standard *in vitro* cell culture is one of the pillars of biomedical science. However, there is increasing evidence that 2D systems provide biological responses that are often in disagreement with *in vivo* observations, partially due to limitations in reproducing the native cellular microenvironment. 3D materials that are able to mimic the native cellular microenvironment to a greater extent tackle these limitations. Here, we report Porous yet Dense (PyD) type I collagen materials obtained by ice-templating followed by topotactic fibrillogenesis. These materials combine extensive macroporosity, favouring the cell migration and nutrients exchange, as well as dense collagen walls, which mimic locally the Extracellular Matrix. When seeded with Normal Human Dermal Fibroblasts (NHDFs), PyD matrices allow for a faster and more extensive colonisation when compared with equivalent Non-Porous matrices. The textural properties of the PyD materials also impact cytoskeletal and nuclear 3D morphometric parameters. Due to the effectiveness in creating a biomimetic 3D environment for NHDFs and the ability to promote cell culture for more than 28 days without subculture, we anticipate that PyD materials could configure an important step towards *in vitro* systems applicable to other cell types and with higher physiological relevance.

## INTRODUCTION

Throughout the last century, traditional *in vitro* culture of adherent cells—the one that is performed in Petri dishes or equivalent 2D rigid substrates—has revolutionized life sciences, from fundamental biology up to routine protocols in clinical practice. Once the minimal physiological conditions are established for a given cell type, traditional *in vitro* cultures allow to scrutinize the cellular response to controlled stimuli, unveiling many physiologic and pathologic processes. Because it reduces biological systems to their simplest forms, cell culture allows to isolate the biological response to a set of chosen parameters, while reducing both the necessary time and resources when compared to *in vivo* systems. Also, from an ethical standpoint, the potential benefits of *in vitro* cell culture experiments are paramount, because it allows to narrow down the conditions to be tested *in vivo*, which, in turn, limits the number of subsequent animal experiments. However, and in spite of these advantages, conventional *in vitro* cell cultures lack the capacity to reproduce the native cellular environment, hindering these systems from responding to complex stimuli as they would *in vivo*. As a consequence, the last decade has witnessed a consensus around the limitations of conventional *in vitro* cell cultures. If these limitations are now widely accepted, the solutions pointed out to tackle them are far more scattered. One of the few common angles resides in the change in dimensionality of the cell culture support. Apart from highly polarized epithelial cells—that naturally occupy 2D environments with variable curvature—most cells *in vivo* are enclosed in a 3D environment. Culturing cells in conditions that mimic the topological aspect of the native environment of cells— 3D cell culture—is thus expected to help bridging the gap between traditional *in vitro* and *in vivo* studies.^1–3^

One central strategy to elaborate physiologically relevant platforms for cell culture is to recapitulate the matrix attributes that favour cellular function in 3D. One of such attributes is porosity. If the pore size, pore shape and pore connectivity are suitable, then porosity determines the pathways for cell migration and, in the absence of vascularization, provides an alternative for the transport of gases, nutrients and cellular debris.^4^ The second attribute depends on the composition of the material. Hydrogels, in particular those prepared from the same macromolecules found in the Extracellular Matrix (ECM), play a critical role in cell culture.^5,6^ Type I collagen—the most abundant protein in connective tissue—is the backbone of the ECM in mammals (bones, skin, cornea, tendon, arteries and veins, etc.) and thus a good candidate to mimic the cellular environment of the ECM in 3D cell culture materials. Beyond its widespread presence in human tissues, type I collagen presents important advantages when compared to other biopolymers; it is non-immunogenic, it remains highly conserved across species, and displays the GFOGER peptide sequence—among others—which enables integrin-mediated cell adhesion.^7^ *In vivo*, type I collagen is found at different hierarchical levels—from the triple α-helices up to the full tissue scale—that contribute to the elastic properties of the tissues and to their physico-chemical stability.^8^ Achieving such degree of hierarchical organisation *in vitro* is a considerable challenge that remains to address. The concentration of collagen in solution, prior to fibrillogenesis, is one of the main levers to access (partially) the degree of order found in dense tissues. When dissolved in acidic aqueous media, type I collagen displays a lyotropic behaviour that ranges from a disordered isotropic phase up to a cholesteric phase. The role of these ordered mesophases in achieving dense fibrillar assemblies after fibrillogenesis has been extensively discussed in the literature and remains a critical parameter in trying to grasp, *in vitro*, the complexity of native tissues.^9^ Despite this knowledge, the concentration of collagen in the elaboration of most biomaterials remains below the specific levels that induce self-organisation. One of the reasons lies in the high viscosity of concentrated collagen solutions, which limits most top down fabrication techniques such as 3D printing^10^ or electrospinning^11^—both heavily dependent on viscosity.

Here we report materials designed to reproduce the complexity of the ECM in 3D cell culture of different tissues, that cumulate the two apparently contradictory notions discussed above: porosity and density. We show that a combination of ice-templating (a technique initially developed for the fabrication of porous ceramics) and topotactic fibrillogenesis^12^ (a strategy that stabilizes frozen collagen monoliths to form self-supported porous fibrillar gels without covalent crosslinking) allows to build materials that harness these two central qualities: Porous yet Dense (PyD). As a direct consequence of the fabrication strategy, these materials combine a macroporous texture and a dense collagen fibrillar architecture, which together yield an appropriate environment for fibroblasts migration and proliferation. The ice-templating process plays a double role: it creates the pores—that correspond to the volume occupied by ice crystals before thawing—and promotes the increased concentration of collagen within the pore walls. We analysed the phase segregation that occurred during freezing and topotactic fibrillogenesis to infer the final local concentration of type I collagen on the matrices’ walls. Fibroblasts were seeded in PyD collagen matrices and either seeded or encapsulated in Non-Porous (NP) matrices generating three different cell culture conditions, PyD, NP-col and NP-enc, respectively. In all three conditions collagen matrices were obtained from equivalent initial concentration (40 mg mL^-1^). The different cell culture systems obtained were compared in terms of the spatial distribution of fibroblasts within the matrices, the cells’ proliferative status, and their 3D morphometric parameters (cytoskeletal and nuclear) during culture up to 28 days without subculture.

## MATERIALS AND METHODS

The chemical reagents used in this work were purchased from Sigma-Aldrich (Merck KGaA, Darmstadt, Germany), unless otherwise stated. The cell culture reagents and supplements were purchased from ThermoFisher Scientific (Gibco™, MA, USA).

### Collagen solution

Type I collagen was extracted and purified from young Wistar or Sprague-Dawley rat tail tendons, under sterile conditions, as previously described.^12^ Briefly, tendons were thoroughly cleaned with phosphate buffered saline (PBS) 1× and 4 M NaCl, then dissolved in 3 mM HCl. Differential precipitation with 300 mM and 600 mM NaCl, followed by redissolution and dialysis in 3 mM HCl, provided a high-purity type I collagen solution, as assessed by SDS-PAGE. Using hydroxyproline titration a concentration of 5.3 ± 0.2 mg mL^−1^ was estimated. This stock solution was stored in aliquots at 4 °C until further use. To obtain solutions of higher concentration (40 mg mL^−1^ and 50 mg mL^−1^), the appropriate volume of stock solution was centrifuged at 3000g at 10 °C in Vivaspin® 20 concentrators (Sartorius Stedim Biotech GmbH, Göttingen, Germany) with a membrane cut-off of 300kDa. Concentrated collagen solutions were also aliquoted in 1 mL syringes (Injekt®-F, B Braun, Hessen, Germany), centrifuged again at 3000g at 10 °C to remove air bubbles, and stored at 4 °C until further use.

### Porous yet Dense (PyD) materials preparation

PyD matrices were produced through coupling of ice-templating technique and collagen self-assembly by a topotactic conversion approach as previously described.^12^ Concentrated collagen solutions (0.1 mL, 40 mg mL^−1^) poured into cylindrical polystyrene moulds (height 10.7 mm, diameter 6.4 mm) were cooled from 20 to −60 °C at a 5 °C min^−1^ rate on a home-made ice-templating setup comprised of an aluminium rod partially immersed in liquid nitrogen, a heating element, a thermocouple to monitor the temperature of the rod’s surface in contact with the solution, and a temperature controller (Omega Engineering Inc, CT, USA). The final temperature of −60 °C was maintained so that samples were entirely frozen. Frozen samples were then stored in a freezer at −80 °C until further use. Collagen self-assembly was carried out as previously reported.^12^ Briefly, frozen samples kept at 0 °C were exposed to ammonia vapours to induce pre-fibrillogenesis. After eliminating the ammonia vapours, the samples were immersed in PBS 5× for 2 weeks in order to complete fibrillogenesis and subsequently stored in sterile DI water at 4 °C until use. The obtained constructs were of 2.5−3.5 mm height and 6.3−6.7 mm diameter.

### Non-Porous (NP) materials preparation

*Non-Porous materials for colonisation (NP-col)*. NP-col matrices were prepared under sterile conditions by dispensing acid-soluble type I collagen (0.1 mL, 40 mg mL^−1^) in 96-well plates, followed by addition of 0.1 mL of Dulbecco’s modified Eagle’s medium (DMEM) on top of the concentrated collagen solution. Plates were incubated for 1h at 37 °C, 5% CO_2_, allowing collagen hydrogels to form. Then, the culture medium was discarded, and the formed hydrogels were further incubated in 0.2 mL of DMEM for 24h. *Non-Porous materials for encapsulation (NP-enc)*. NP-enc gels (0.1 mL, 40 mg mL^-1^) were prepared on ice by mixing in the wells ice-cooled collagen solution (80 µL, 50 mg mL^−1^), PBS 1× (10 µl), NaOH 0,1 M (2.4 µL) in order to neutralise the collagen solution, and complete DMEM (7.6 µL), alone or with the appropriate number of cells, as described below. Collagen gels were completely set after the addition of 0.2 mL DMEM and subsequent plate incubation for 1h at 37 °C, 5% CO_2_.

### Cell culture

Normal human dermal fibroblasts (NHDFs, PromoCell, Heidelberg, Germany) at passage 7 were cultured in low-glucose DMEM supplemented with 10% foetal bovine serum (FBS) and 1% penicillin/streptomycin, at 37 °C, 5% CO_2_, in a humidified atmosphere. When reaching confluence, NHDFs were passaged by treatment with 0.1% trypsin and 0.02% EDTA, resuspension in the above culture medium, and seeding in new flasks at adequate numbers. The fibroblasts were counted with a haemocytometer and their viability was assessed by trypan blue exclusion.

### 3D cell culture

*PyD samples*. The day before cells seeding, already prepared PyD matrices were rinsed 2-3 times in cell culture medium for 24h. On day 0, PyD matrices were transferred into wells of 96-well plates oriented so that pore openings were facing the top of the well, and 10 000 NHDFs per well were seeded on top of the materials. *NP-col samples*. Samples were cellularised similarly to porous materials; 10 000 NHDFs were deposited onto the upper surface of the collagen cylinders. *NP-enc samples*. Cells encapsulation was performed by mixing complete DMEM (7.6 μL) with 10 000 suspended NHDFs during the NP-enc materials preparation, as described above. Seeded and encapsulated cells were cultured in triplicates for 28 days in 0.2 mL complete DMEM per well, changing the culture medium every other day. Negative controls were also prepared for each matrix type.

### Scanning Electron Microscopy (SEM)

Structural features of all three types of materials were firstly observed on lyophilised dry samples, coated with a 10 nm gold layer, under a Hitachi S-3400N microscope (operating at 3 kV). Pores size of the PyD samples was determined using automated image analysis with FIJI software^13^ (see Supplementary Information) on binarized images of the samples surface. The characteristic dimensions of the pores were calculated based on at least 650 different pores using minimum Feret’s diameter (minimum calliper) measurements of individual pores. Fibrillar hydrated materials were also characterised after an overnight fixation step in paraformaldehyde (PFA) 4% w/v in PBS 1× at 4 °C, followed by dehydration in serial ethanol baths of increasing concentrations, and supercritical drying (Leica EM CPD300, Leica Microsystems, Wetzlar, Germany).

### Transmission Electron Microscopy (TEM)

Hydrated samples were fixed overnight with PFA 4%, rinsed with sodium cacodylate buffer 0.1 M and saccharose 0.6 M, further fixed with glutaraldehyde 8% for 2h at 4 °C and then Nuclear Fast Red 0.1% for 1h at room temperature. Samples were subsequently dehydrated in serial ethanol baths of increasing concentrations, progressively transferred to propylene oxide, and incorporated in araldite resin (Electron Microscopy Sciences, PA, USA) prior to sectioning (Leica EM FC7, Leica Microsystems). Then, ultrastructural collagen features were observed on 70 nm ultrathin sections contrasted with uranyl acetate, under a transmission electron microscope (Tecnai Spirit G2, FEI Company, Thermo Fisher Scientific, MA, USA) operating at 120 kV, equipped with an Orius™ SC1000 CCD camera (Gatan, CA, USA).

### Histological sections analysis

Hydrated samples were fixed overnight with PFA 4%, then dehydrated firstly in serial ethanol baths of increasing concentrations and then in butanol-1. Dehydrated samples were embedded in paraffin and then cut in 7 μm thick sections using a rotary microtome (HistoCore AUTOCUT, Leica Biosystems, Nussloch, Germany). Sections were deparaffinised, rehydrated, stained with picrosirius red or haematoxylin, and dehydrated again using ethanol and toluene. Before observation, stained sections were mounted between glass slide and coverslip using the Eukitt® Quick-hardening mounting medium. Slides were observed under an Eclipse E600Pol microscope (Nikon, Tokyo, Japan) equipped with a DS-Ri1 camera (Nikon). Images were also acquired under built-in cross-polarisers to probe the birefringence of the samples.

### Fluorescent labelling

At days 1, 14, and 28 of cell culture, the corresponding PyD, NP-col, and NP-enc samples were rinsed with PBS 1× and fixed overnight in 4% PFA. Longitudinal sections (cut along the cylinders’ axis) of 250 μm thickness were obtained from agarose-embedded fixed samples using a vibration microtome (Compresstome® VF-300-0Z, Precisionary Instruments, NC, USA), with a double-edged stainless-steel blade oscillating at 10 Hz at a cutting speed of 3-4 mm s^−1^. All obtained slices were stored in PBS 1× at 4 °C until the fluorescent labelling procedure.

Cells in all three types of constructs were labelled with TO-PRO™-3 (Invitrogen™, ThermoFisher Scientific) for the nuclei, Alexa Fluor™ 488 phalloidin (Invitrogen™) for the F-actin filaments, and anti-Ki67 primary antibody (ab 15580, Abcam, Cambridge, UK) followed by Alexa Fluor™ 555 donkey anti-rabbit secondary antibody (A-31572, Invitrogen™) to study the cells proliferation. The slices were firstly observed under an Axio Imager.D1 fluorescence microscope (Zeiss, Oberkochen, Germany) to confirm cells migration and ensure the absence of any contamination. Representative slices for each construct type and culture time point were selected for observation under a confocal microscope.

### Confocal and multiphoton microscopy

Selected slices were further observed under a Leica DMI6000 Upright TCS SP5 confocal microscope (Leica Microsystems) coupled with a femtosecond Ti:Sapphire laser (Mai-Tai® DeepSee™, Spectra-Physics, CA, USA), thus enabling the combination of fluorescence and second harmonic generation (SHG) observations. Z-stacks and tile scans in representative volumes (40-60 μm^3^) and areas of the samples were acquired using a 25× water immersion objective (numerical aperture 0.95, working distance 2.5 mm, Leica Microsystems). Fluorescence emitted from TO-PRO™-3 and Alexa Fluor™ 488 labelling was detected using two internal hybrid detectors, while Alexa Fluor™ 555 signal was detected by a photomultiplier tube (PMT). SHG imaging of type I collagen was performed by an external hybrid NDD (non-descanned detection) detector as well.

### Image analysis

Image processing and analysis to convert microscopy images into quantitative data were performed using the open-source package FIJI.^13^ Briefly, particle analysis was performed on SEM images to determine the pores size of PyD matrices. Following the appropriate treatment of z-stacks obtained with confocal microscopy, two-dimensional image segmentation was used to evaluate the migration and proliferative activity of NHDFs in the constructs. The minimum distance for all thereby defined cells compared to the reference line of NP-col and PyD matrices surface was also determined. Cells counting and morphometric analysis were performed after the three-dimensional segmentation of z-stacks. Finally, colocalization analysis of TO-PRO™-3 and Ki67 labelling was conducted. A detailed description of acquired z-stacks processing as well as of the macroinstructions and plugins used in this work can be found in Supplementary Information.

### Data plotting and statistical analysis

The graphs and the statistical analysis presented in this work were performed using the scientific applications QtiPlot (version 1.0.0-rc15, IONDEV SRL, Bucharest, Romania) and Prism (version 7.0, GraphPad, CA, USA). The normal distribution of the data was assessed using the Shapiro-Wilk test. Significant differences between multiple groups were evaluated using non-parametric Kruskal-Wallis analysis of variance (ANOVA) followed by uncorrected Dunn’s post hoc (α = 0.05). Differences were considered significant for *p < 0.0332, **p < 0.0021, ***p < 0.0002, ****p< 0.0001.

## RESULTS AND DISCUSSION

### Materials for 3D cell culture

Using ice-templating as a strategy to tailor biomaterials has become increasingly popular in recent years due to the intrinsically mild conditions involved in the process, ensuring biomolecules are not denatured, in combination with the ability to tailor the materials’ macroporosity using simple parameters such as the ice front velocity or the processing temperatures.^14–16^ To determine the relevance of the materials prepared by this technique as 3D cell culture platforms we have elaborated two types of matrices from 40 mg mL^-1^ type I collagen solutions. PyD matrices were prepared by ice-templating followed by topotactic fibrillogenesis (Fig. 1A, left) whereas NP matrices were obtained by incubating a collagen solution in presence of DMEM to induce gelation (Fig. 1A, right). The resulting structures are depicted by the 3D reconstructions of a strip of each matrix acquired by confocal microscopy from SHG signal (Fig. 1B). Only co-aligned collagen triple helices generate SHG signal, due to the formation of non-centrosymmetric structures.^17^ The PyD matrix (Fig. 1B, left) features highly aligned collagen zones (in grey) throughout the whole sample separated by zones where no signal is detected (in white). The latter could be ascribed to pores generated during the PyD fabrication process. The NP matrix (Fig. 1B, right) featured a continuous SHG signal without pores in the length scale observed by confocal microscopy. For both samples the green surface indicates the cell seeding zone used in the subsequent 3D cell culture experiments. Thin sections of the two matrices observed between cross polarisers, under the optical microscope, presented radically different motifs (Fig. 1C, top row). In PyD matrices the birefringence observed by CPLOM (Cross Polarised Light Optical Microscopy) within the pore walls revealed the highly organised state of collagen throughout the material.^9,18^ It also revealed domains of different molecular orientations from one another changing in more or less continuous rotation (See Fig. S1 in Supplementary Information for more exhaustive characterisation). The increase of collagen concentration during the freezing process forces the system from isotropic to anisotropic organisation due to the natural lyotropic mesogen nature of collagen in acidic condition.^9^ These dark zones were surrounded by interstices—the pore walls—with alternating extinction and bright zones, characteristic of dense ordered gels (Fig. 1C, top left, Fig. S1). These results confirm the formation of fibrillar ordered domains, most likely originating from fibrillogenesis of collagen molecules pre-aligned during the freezing step. On the contrary, NP samples (Fig. 1C, top right) did not present bright zones under the CPLOM irrespective of the angle between the sample and the cross polarisers. The absence of birefringence indicates an isotropic arrangement of collagen within the samples. These results were comforted by the observation of both types of samples under the SEM (Fig. 1C, middle row). PyD samples display alternating areas of collagen fibrils bundled together into dense zones and sparse zones where loose fibrils are visible. Contrary to PyD samples, NP materials presented an isotropic distribution of collagen fibrils, without any visible spatial variation in terms of their packing density, in line with observation made by SHG and CPLOM. Ultrathin samples of both materials inspected under TEM allowed to grasp further details about the local arrangement of collagen fibrils for both types of materials (Fig. 1C, bottom row). At the suprafibrillar level, PyD samples are structured as densely packed collagen fibrils where large alignment domains coexist with twisted arrangements. In the immediacy of these motifs, a pore (Fig. 1C, bottom left, upper corner of the image) shows no collagen motifs. NP samples were composed of randomly oriented collagen fibrils scattered throughout the ultrathin section, which is in good agreement with a lower local concentration in these materials and with the result obtained by the other microscopy techniques. Ice-templating followed by topotactic conversion favours both the formation of a porous structure and the local concentration of collagen molecules. The combination of these factors translates into ordered fibrillar domains after gelling: porous, yet dense!

**Figure 1.**
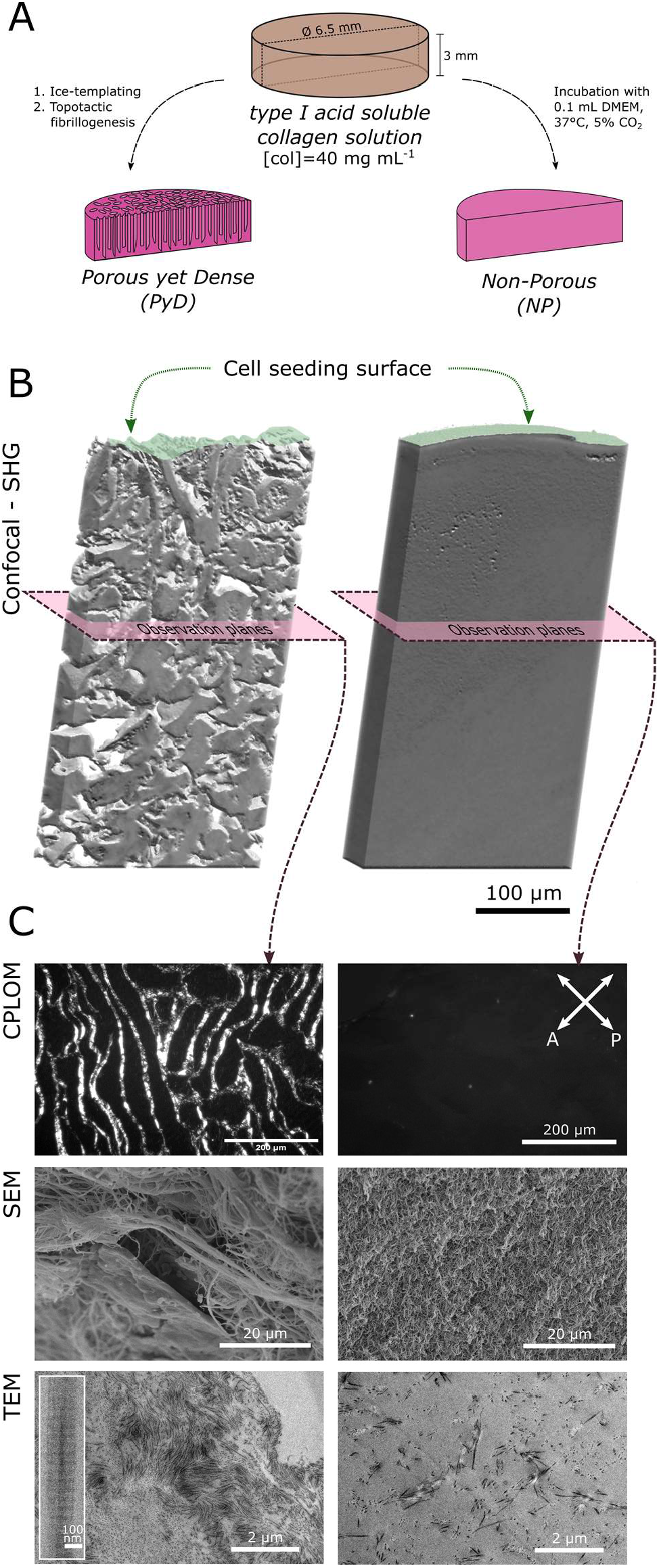
Multiscale characterisation of collagen matrices. A) Simplified fabrication scheme for Porous yet Dense (PyD) and Non-Porous (NP) samples. B) 3D representation of confocal microscopy images of PyD and NP matrices obtained from SHG signal. Green surface corresponds to cell seeding zone in the subsequent cellularisation studies. Pink plane illustrates the approximate orientation and position of the following observations under the microscope. C) Microscopic characterisation of the collagen matrices using CPLOM (top row), SEM (middle row), and TEM (bottom row). Inset in TEM image of PyD sample depicts an individual collagen fibril at higher magnification.

The pores in between the dense collagen walls, accessible from the sample surface (Fig. S2), ranged in size (minimum Feret diameter) from 6 to 198 µm with an average value of 20±16 µm. Their connection between the sample surface and the material inner porosity could act as the predominant migration pathways during cell colonisation, but also act as privileged exchange routes for liquid transport and gas diffusion.

### Collagen local concentration in PyD materials

As discussed above, increasing the concentration of collagen is one of the main levers to partially mimic the ordered nature of the ECM. Analysis of the volume fraction of PyD collagen matrices using Trainable WEKA Segmentation^19^ on histological section of the PyD matrices stained with haematoxylin, allowed to infer the local concentration of the collagen within the materials pore walls. Before freezing, collagen (40 mg mL^-1^) was equally distributed within the solution. Upon freezing, pure ice crystals were formed, leading to the concentration of collagen within the interstices corresponding to the pore walls. These interstices were subsequently transformed by topotactic conversion into fibrillar collagen domains. Since the samples did not experience any macroscopic volume variation upon fibrillogenesis, it was possible to establish the final local concentration of collagen in the matrix walls. The surface fraction of the histology section imaged under the optical microscope (OM) is presented in Fig. 2 left, and the corresponding segmented image in Fig. 2 right. The area fraction of pore walls was *ca* 0.3, which corresponded to a local collagen concentration of *ca*. 130 mg mL^-1^. This value lies well above the isotropic/anisotropic critical concentration of collagen acidic solutions,^20^ and explains the birefringence observed in CPLOM images. This quantification demonstrated that the local concentration of type I collagen in PyD matrices approached those found in the ECM of soft native tissues.

**Figure 2.**
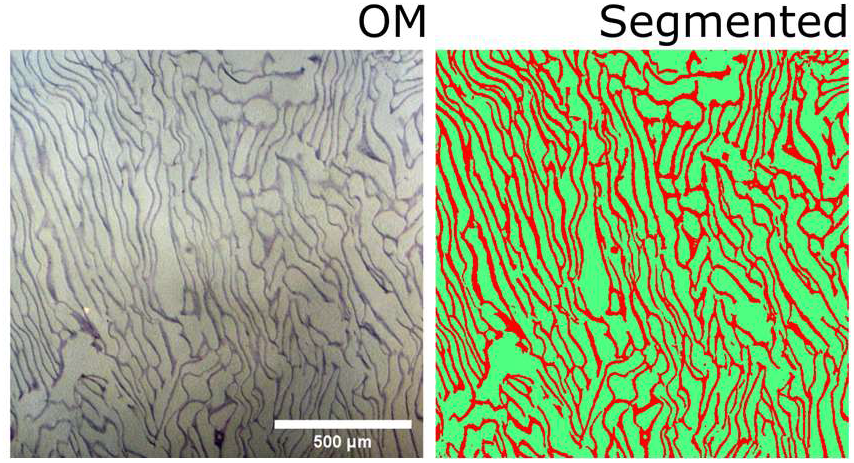
Histological section of PyD matrices allows to determine the local concentration of type I collagen. Left: Optical microscopy of a histology section of 7 μm PyD matrix stained with haematoxylin. Right: Segmented image corresponding to the collagen walls fraction (red) and pores (green).

### Colonisation of the collagen-based materials by seeded fibroblasts

NHDFs were seeded on PyD and NP-col materials (Fig. 3 Day 0) and the colonisation occurring from the top of the substrates was studied over time. At day 1, the position of all NHDFs nuclei was located at the seeding surface. At day 4, fibroblasts seeded on the PyD matrices had already started migration into the porous matrices (data not shown) whereas cells were still at the surface of NP-col matrices. Cells migration and colonisation inside the PyD matrices extended over a distance up to 260 µm at day 14 and up to 750 µm from the seeding surface at day 28 (Fig 3. Days 1, 14, and 28), which corresponds to a continuous migration rate over time. NHDFs migration into the NP-col matrices only started after the first two weeks of culture, in complete agreement with previous results reported in the literature.^2,21^ At day 28 the maximal distance attained by the cells was around 110 µm (Fig. 3 Days 1, 14, and 28), which corresponds to a migration rate at least six times inferior when compared to the PyD matrices. These results highlight the key role of the controlled porosity induced by ice-templating, which triggers a rapid and extended colonisation of the sample volume by NHDFs, whereas NP-col collagen gels induced 2D-like colonisation up to day 14 and limited 3D colonisation at day 28. Such differences could result from the fundamentally different migration mechanisms involved according to each sample texture. The porous network in PyD samples provides a support compatible with amoeboid migration whereas NP samples require protease-dependent migration. The consequences in terms of the application of these matrices as 3D cell culture matrices are most relevant. Porous matrices (PyD) configure a solution to rapidly expose NHDFs to a real 3D environment whereas waiting for over 15 days of cell culture to have cells in an actual 3D environment (NP-col) is highly constraining. Another important point that emanates from the colonisation experiments concerns the absence of passage. NHDFs cultured in standard 2D *in vitro* systems usually require frequent culture medium change and subculture, every 2 and 4-5 days, respectively. Here, NHDFs were cultured up to 28 days without subculturing. Extension of the culture time to 58 days proved possible and coherent with the absence of noticeable sample volume retraction (data not shown). Removing the subculture step from cell culture eliminates the need for proteolytic dissociation procedures such as trypsinisation^22,23^ or other cell detachment techniques^24^ that are required in 2D cell cultures. Being able to maintain long term cell culture without subculturing steps is a remarkable advantage that strengthens the physiological relevance of 3D cell culture and reduces the overall cost of the process. To the abovementioned results add the dramatic differences in number of nuclei detected in the samples at different time points. These are not strictly quantitative results *per se* but provide a strong indication of how the cell proliferation is modulated by the texture of the collagen matrices.

**Figure 3.**
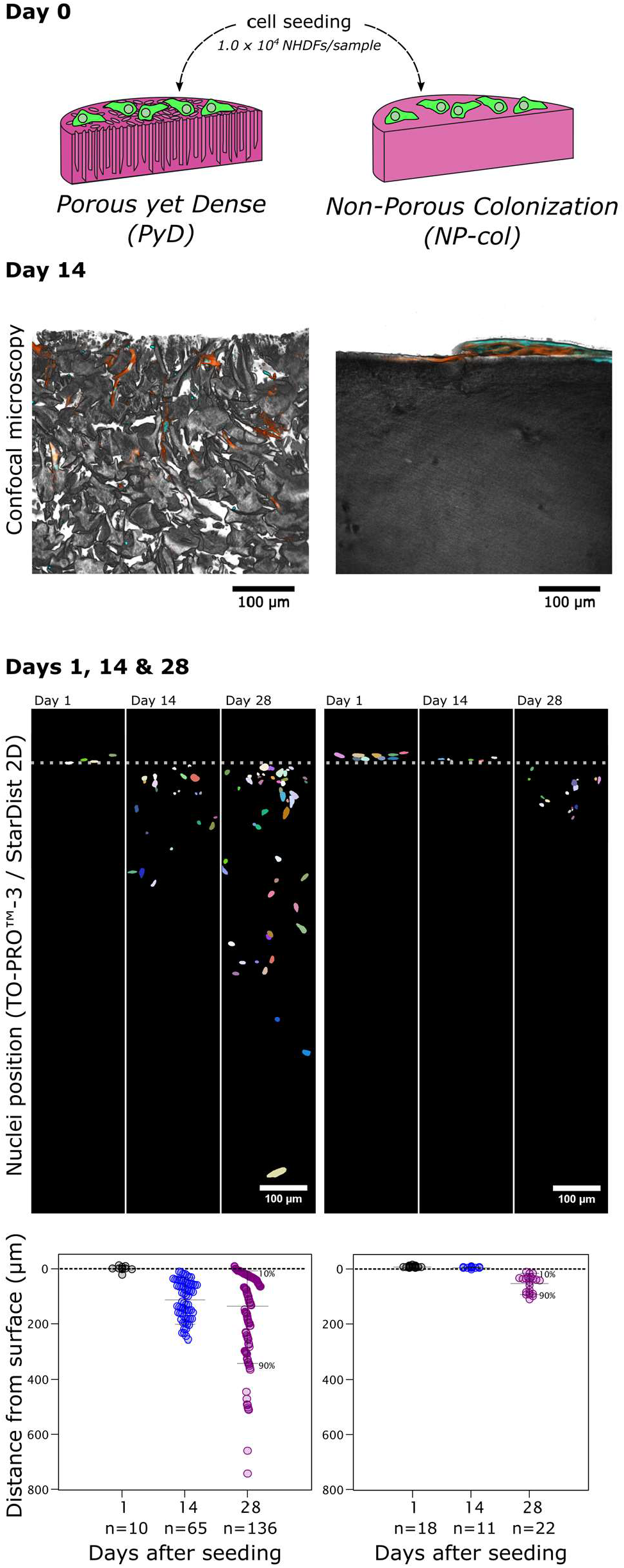
Cell colonisation of PyD and NP materials. Day 0: Scheme of NHDF cell seeding on top of the two different types of substrates. Day 14: Snapshots from 3D reconstructions of confocal SHG/fluorescence images showing NHDFs colonisation inside both types of constructs at day 14. Planes of observation are along the cylinders’ axes and the surfaces in contact with the cell culture medium are on top of the images. Inverted grayscale: collagen (SHG signal), orange: actin (phalloidin). cyan: nuclei (TO-PRO™-3). Days 1. 14 and 28: top) Representative regions of interest (ROI) of processed output images of the StarDist plugin used on Z projections of confocal microscopy images (TO-PRO™-3 staining) showing the evolution over time of nuclei positions. The dotted line represents schematically the top surface of the matrices, bottom) Scatter plots of the calculated minimum distances of each considered nucleus centroid to the reference matrix surface.

### Proliferative status of seeded/encapsulated fibroblasts and colocalization analysis

The proliferative state of NHDFs was assessed based on Ki67 immunolabelling. The corresponding fluorescent signal indicated a continued proliferation in cells in the different samples examined, throughout the cell culture time period.

Colocalization analysis of TO-PRO™-3 and Ki67 labelling in fibroblasts nuclei—conducted using the Colocalization Colormap plugin^25^ (see Supplementary Information)—further confirmed that the cells inside the matrices maintained their proliferative state over time, irrespective of the matrices’ texture. More specifically, the colocalization index values obtained in both types of samples ranged from 0.4778 to 0.8065, indicating a non-random colocalization of signals (Fig. 4A, Fig. S3, Table S1). Of note, in PyD and NP-col matrices, proliferating cells were identified all along the migration pathway in each sample up to the maximal distances attained, hence showing that porosity—obtained either by the ice-templating process in case of PyD samples, or matrix remodelling in the case of NP-col samples—ensures the efficient diffusion of nutrients throughout the whole sample volume. This is consistent with the observation of a growing number of cells inside the matrices, in particular for PyD samples. The colocalization index values of cells in NP-enc samples revealed a rather steady proliferation status as well (Table S1, Fig. S3). However, while maintaining this state, important morphologic alterations in encapsulated cells occurred, as described below.

**Figure 4.**
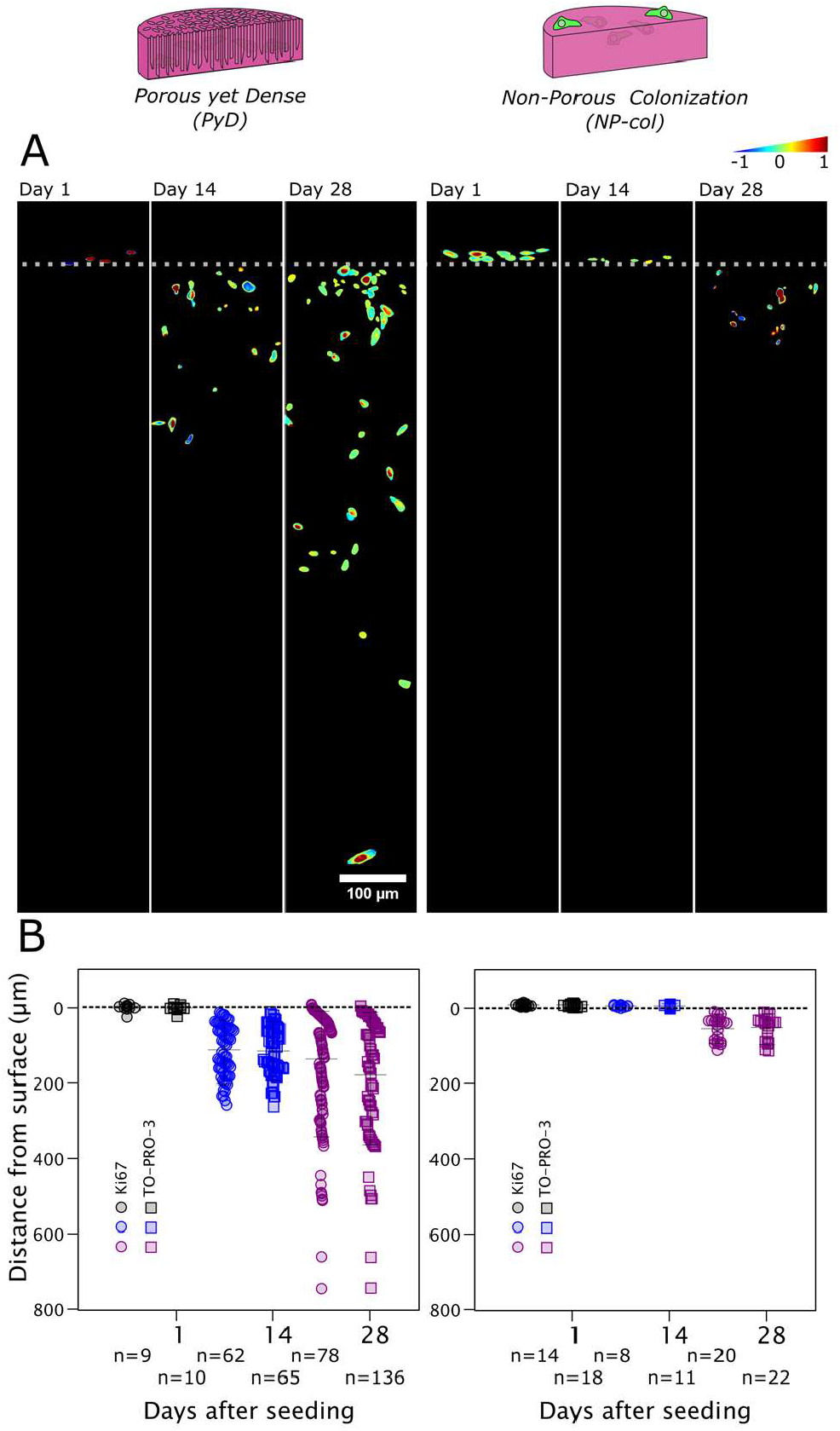
Proliferative status of colonising cells. A) ROI of colormaps on Z projections at days 1, 14, and 28 of confocal microscopy images with colocalization analysis of TO-PRO™-3 and Ki67 labelling. Colocalization indexes range from −1 (blue), corresponding to no-colocalization, to 1 (red), corresponding to absolute colocalization of pairs of corresponding pixels (see the Colormap scale). The dotted line represents schematically the top surface of the matrices. B) Scatter plots of the calculated minimum distances of each considered nucleus centroid to the reference matrix surface for both TO-PRO™-3 and Ki67 labelling (see Fig. S3 and Table S1 for more details).

### Morphometric analysis of the seeded/encapsulated fibroblasts

The quantification of NHDFs 3D morphological features was conducted at both the cytoskeletal and the nuclear level to explore the impact of the porous or non-porous microenvironment on their spatial configurations over time. The data are reported as mean ± SD values.

The cellular architecture of fibroblasts was first assessed based on the cell *volume* (Fig. 5B) and the cell *surface to volume ratio* (S/V, Fig. 5C) as 3D morphometric descriptors, measured based on the actin labelling. Fibroblasts seeded on 2D-like surfaces presented a flat, bi- or multipolar, spindle-shaped morphology. On the contrary, NHDFs featured an expanded bi- or multipolar phenotype with cytoskeletal protrusions in their surrounding 3D microenvironment when migrating into the matrices (Fig. S4). In line with these broad morphological observations, the cell volume tended to increase over time (day 1: n=2, 580 and 1329 μm^3^, day 14: 1979 ± 1556 μm^3^, and day 28: 3409 ± 2861 μm^3^) for fibroblasts seeded on PyD matrices. Moreover, the same cells adopted a more irregular shape in the matrices and seemed to present lower S/V values over time (day 1: n=2, 2.23 and 3.69 μm^-1^, day 14: 1.85 ± 0.52 μm^-1^, and day 28: 2.09 ± 0.78 μm^-1^). Increasing collagen concentration has been reported as a central factor in reducing O_2_ diffusion coefficient in non-porous samples with deleterious consequences to cell viability.^26^ The S/V values for the fibroblasts located in the pores may suggest an efficient O_2_ diffusion without substantial alterations to the obtained cellular morphology.

**Figure 5.**
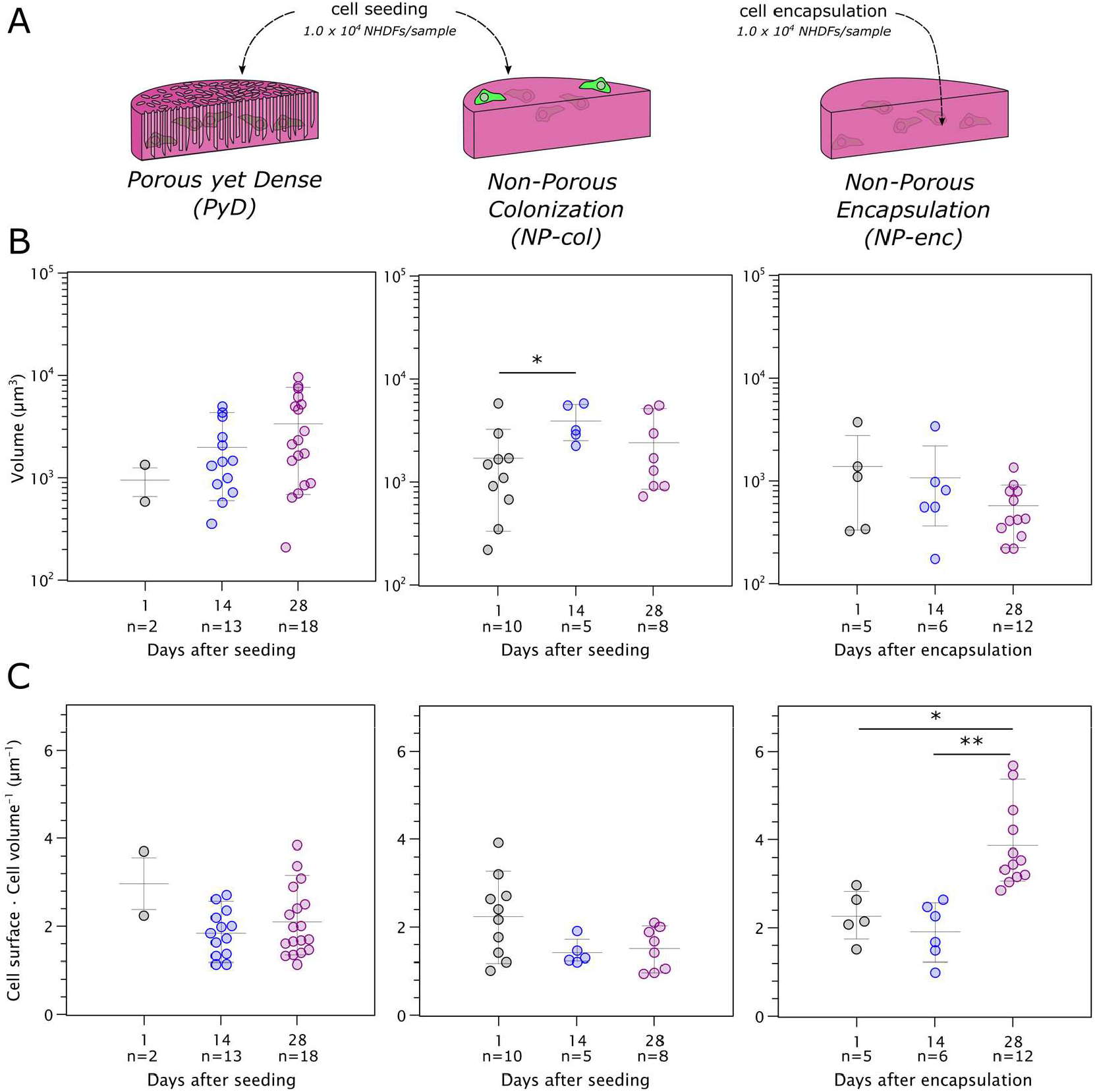
3D morphometric analysis at the cytoskeletal level of fibroblasts colonising PyD and NP collagen materials, based on actin labelling. A) Schematic representation of cell seeding on PyD and NP-col and cell encapsulation in NP-enc matrices. Scatter plots (mean ± interdecile range plotted) of B) the cellular volume and C) cell surface / cell volume ratio measurements of NHDFs nuclei at days 1, 14, and 28 after seeding. Statistically significant differences between different time points for the same material are shown (*p < 0.0332, **p < 0.0021).

Cytoskeletal morphology modifications were also observed for fibroblasts seeded on NP-col collagen gels. The volume of NHDFs at day 1 (1696 ± 1660 μm^3^) was lower than that at day 14 (3977 ± 1663 μm^3^, p = 0.0371) and day 28 (2393 ± 1939 μm^3^). The cells also exhibited a heterogeneity in S/V (2.243 ± 0.919 μm^-1^), which was lower at days 14 (1.422 ± 0.289 μm^-1^) and 28 (1.506 ± 0.476 μm^-1^). These trends are consistent with the previously described behaviour of fibroblasts seeded on dense collagen gels.^21^ Proliferating NHDFs increased their individual cellular volume up to the confluence level at day 14 and then started collagen gels remodelling and colonisation, maintaining a larger cellular volume than at their initial elongated and flatter state.

Fibroblasts embedded in the collagen gels (NP-enc) did not maintain their cellular volume homeostasis, and a decreasing trend in their volume over time was found (day 1: 1379 ± 1397 μm^3^, day 14: 1084 ± 1178 μm^3^, and day 28: 571 ± 342 μm^3^). In addition, a dramatic change was observed for the S/V at day 28 (3.857 ± 0.947 μm^-1^) compared to days 1 (2.269 ± 0.561 μm^-1^, p = 0.0368) and 14 (1.920 ± 0.641 μm^-1^, p = 0.0018). Encapsulated cells in concentrated collagen matrices (40 mg mL^-1^) are unable to remodel their microenvironment so they are rather found increasingly confined and “immobilized” in their embedding material.^4,27^ The combination of decreased cellular volume and high S/V at day 28 suggests that nutrients and waste diffusion in and out of fibroblasts were inhibited in the NP-enc matrices leading to reduced cellular metabolism. The S/V value for fibroblasts in NP-enc matrices was also significantly different from those obtained at day 28 in PyD (p = 0.0001) and NP-col (p < 0.0001) matrices (Fig. S5B), further highlighting the importance of the pore network for an efficient transport of materials between the cell and the cell culture media. Moreover, based on the extensive non-specific signal obtained at day 28 in NP-enc matrices (at the nuclear level as well), the presence of cellular debris and fragments is highly possible, suggesting that part of the cells could be in an apoptotic state.

The cellular architecture of fibroblasts was further assessed at the nuclear level using the *ellipsoid flatness* (EF, Fig. 6A) and the (corrected) *sphericity* (Sph, Fig. 6B) measurements of the nuclei, which estimate the compression along their diameter to form an ellipsoid (lower EF values indicate a more spherical shape) and the resemblance of their shape to the hypothetical nuclear morphology of the perfect sphere (where Sph=1), respectively.^28^ As described above, fibroblasts on 2D-like surfaces presented a spindle-shaped morphology, whereas during cells migration through the pores of the PyD materials a shift from their spindle-like to a larger and stellate configuration was observed. The nuclei of fibroblasts situated on the samples surface were characterised by an elongated shape, namely on PyD matrices at day 1 (EF 1.89 ± 0.38) and NP-col matrices at days 1 (EF 1.74 ± 0.52) and 14 (EF 1.65 ± 0.48) after seeding. Besides, an ellipsoid shape was found for the nuclei of cells that had migrated into the matrices, namely in PyD at days 14 (EF 1.42 ± 0.26) and 28 (EF 1.42 ± 0.27) and NP-col at day 28 (EF 1.49 ± 0.26). Moreover, EF values of NHDFs nuclei seeded on PyD matrices were significantly higher at day 1 compared to the values at days 14 (p = 0.0149) and 28 (p = 0.0150). A similar trend was observed for the Sph as the nuclei presented at day 1 higher values (0.74 ± 0.03) compared to days 14 (0.65 ± 0.06, p = 0,0212) and 28 (0.68 ± 0.10). These findings are consistent with the cells 3D distribution and shape adaptation: at day 1 the fibroblasts on the samples surface had a 2D-like morphology whereas at days 14 and 28, as cells migrated into the constructs moving through the pores to occupy the available space, their nuclei partially lost their spherical shape. The opposite trend was observed for the Sph of NHDFs nuclei seeded on the NP-col collagen hydrogels. More specifically, the values at day 1 (0.63 ± 0.08) were the lowest compared to these at day 14 (0.69 ± 0.10) and day 28 (0.73 ± 0.06, p = 0,0051). Previous studies have shown that fibroblasts seeded on dense collagen matrices grow as a monolayer until they attain confluence at day 14, and then they migrate into the gels through matrix remodelling.^2,21^ During the enzymatic digestion of the matrix— occurring in NP-col samples—their shape is not subjected to the alterations occurring during migration through pre-formed pores of the macroporous structure—occurring in PyD samples—, and the nuclei maintain a more spherical shape.^27^

**Figure 6.**
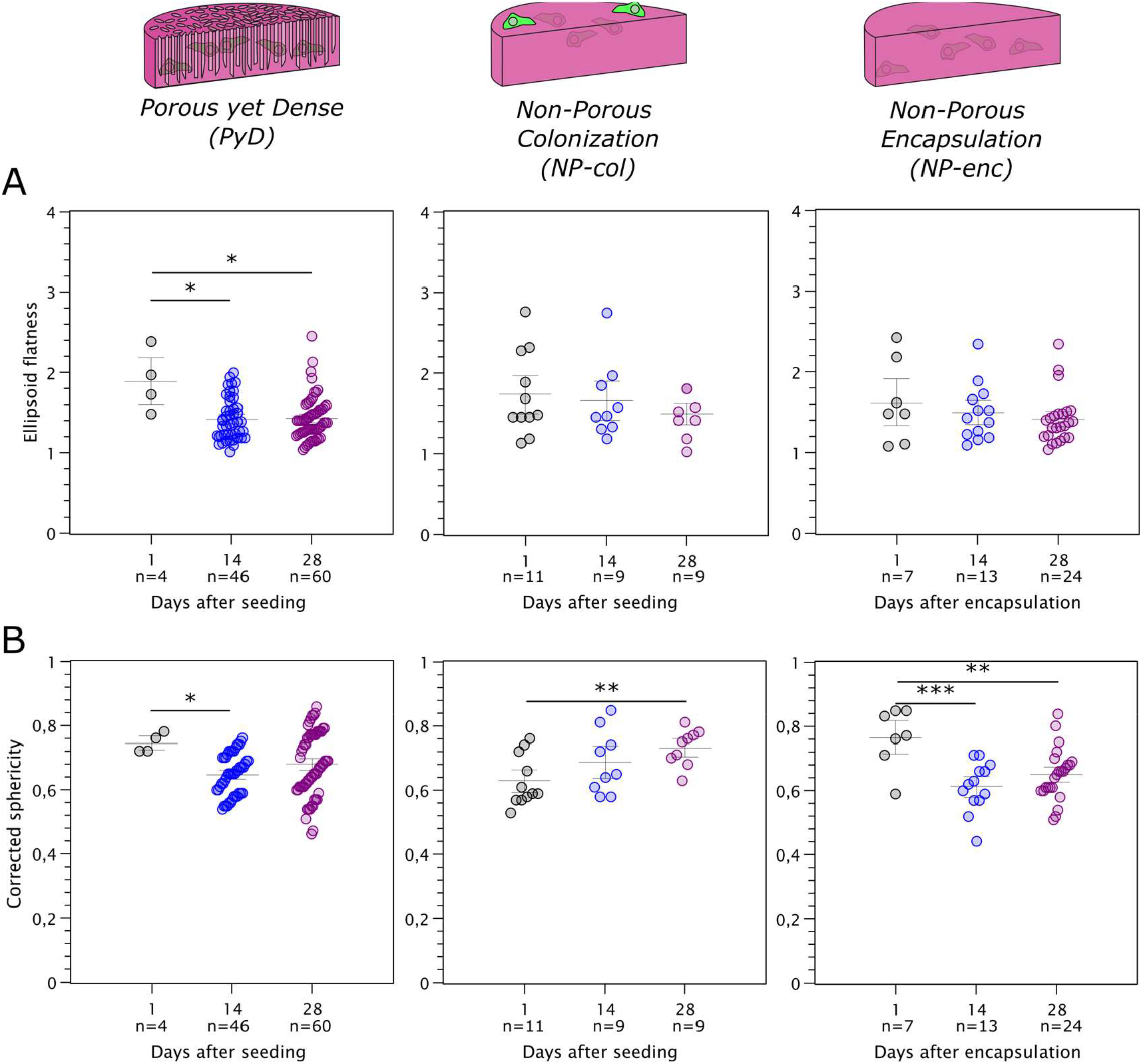
3D morphometric analysis at the nuclear level of fibroblasts colonising PyD and NP collagen materials, based on TO-PRO™-3 labelling. Scatter plots (mean ± interdecile range plotted) of: A) the ellipsoid flatness and B) corrected sphericity measurements of NHDFs nuclei at days 1, 14, and 28 after seeding. Statistically significant differences between different time points for the same material are shown (*p < 0.0332, **p < 0.0021, ***p < 0.0002).

Fibroblasts in NP-enc hydrogels acquired an overall convex configuration without protrusions, maintained throughout the cell culture duration. The nuclei of encapsulated NHDFs at day 1 exhibited an elongated configuration (EF 1.62 ± 0.51), and at days 14 and 28 an ellipsoid one (EF 1.49 ± 0.35 and 1.41 ± 0.31, respectively). In terms of sphericity, the nuclei of NHDFs embedded in the collagen gel had a significantly higher mean value (0.77 ± 0.09) at day 1 compared to days 14 (0.61 ± 0.08, p = 0.0010) and 28 (0.65 ± 0.08, p = 0.0062). Thus, nuclei shape modifications followed a trend corresponding to a more “3D matrix-like” environment, similar to the one observed in PyD matrices.

This is also highlighted by the differences observed between the Sph values of nuclei at days 1 and 28 for NP-col versus NP-enc matrices (p = 0.0026 and p = 0.0123, respectively, Fig. S6).

These results illustrate the importance of the topology of the 3D cell culture matrices on the morphometric parameters of the cells. Beyond the expected changes in the morphometric parameters of the cytoplasm, whose shape directly depends on the adhesion between the cells and the surrounding matrices, we observed important 3D morphometric changes at the nuclear level.

## CONCLUSIONS

The expectations that 3D cell culture would bridge the gap between standard *in vitro* and *in vivo* studies have continuously risen in the past decades. However, the quest to design materials that can mimic the ECM in long-term cultures and that can be extensively colonised is still ongoing. Porous yet Dense matrices introduced here display a local concentration of type I collagen that rises from initial 40 mg mL^-1^ to *ca*. 130 mg mL^-1^, reaching local values that are in line with the concentrations of collagen in native tissues.^29^ Because of the lyotropic behaviour of collagen in solution, this change in local concentration translates into highly organised collagen domains, which reproduce a part of the multi-scale hierarchical order of collagen in the ECM. These characteristics allow to tackle part of limitations found in current materials designed for 3D cell culture. Analysed up to 28 days, PyD materials induce radically different migration kinetics and cell density as compared to equivalent non-porous matrices—same composition and average collagen concentration. Because of the predefined porosity, cells seeded on PyD matrices start migration from day 4, whereas cells seeded in equivalent non-porous matrices did not start to migrate before day 14. This difference in the kinetics of migration implies dramatic gains in terms of the efficiency of a 3D cell culture model since the required time for cells to be in an effective 3D environment is shortened by 10 days.

Analysis of the 3D morphometric parameters of the cytoplasm of the seeded fibroblasts has shown that the topological differences of the collagen matrices have a direct impact on the volumes and surface to volume ratios of the cells. These differences were found to extend down to the nuclear morphometric parameters, which highlights the role of the textural properties of the matrices in modulating cell behaviour.

In summary, the PyD materials presented here provide an interesting alternative to the current 3D cell culture platforms. The effective colonisation by adherent cells without additional biochemical signalling, the ability to perform long-term cell culture experiments without the disruption induced by cell passaging and the enhanced physiological relevance of the PyD matrices configure strong arguments to replace the current solutions for 3D cell culture.

## Author Contributions

The manuscript was written through contributions of all authors. All authors have given approval to the final version of the manuscript.

## Acknowledgments

The authors would like to thank Bernard Haye and Corinne Illoul for their excellent technical assistance in samples preparation for TEM and histological observations respectively. The authors also thank Julien Dumont (Centre for Interdisciplinary Research in Biology, Collège de France – CIRB, CdF) for his precious help in image acquisitions using confocal fluorescence and SHG microscopy, and Florian Fage for his help in image analysis. This work has been supported by the Île-de-France Region through RESPORE (Network of Excellence in Porous Solids).

## Supplementary information

### Materials and Methods

#### Image analysis

For all acquired z-stacks and tile scans, the channels were split and appropriate regions of interest (ROI) were defined for the subsequent analyses. Standard processing and noise reduction treatment (Despeckle, Remove outliers, Subtract background, Thresholding) and filtering were used based on the intended analysis.

#### Pore size measurements (Analyze Particles)

SEM images of porous scaffolds surfaces were binarized after being processed with threshold and noise removal. Particle analysis was used to measure Feret’s diameter of the pores in each image. Pores size was determined through MinFeret values, the minimum calliper diameter.

#### Nuclei position (TO-PRO™-3 staining) – 2D (StarDist plugin)

Before applying a Z-projection on the TO-PRO™-3 channel, the z-stack was pre-processed for noise reduction followed by Thresholding. The StarDist plugin^1^ was applied on this Z-projection, set on Versatile (fluorescent nuclei) model and on default parameters to normalise the image. Several trials were performed adjusting the NMS (non-maximum suppression) settings so to obtain the best possible segmentation for the objects (nuclei). The output images were cross-checked with the z-stacks for nuclei and F-actin labelling to exclude artefacts. The selected produced image was further treated with the 3D manager e.g., to erase objects that corresponded to artefacts. The ROI set produced by the StarDist plugin was corrected based on the final image and measurements of the nuclei were performed.

#### Minimum distance: cells-scaffold’s surface – 2D (Macro Reference Distances)

Using this macro^2^ the minimum (and maximum) distances for all objects compared to a reference file of XY coordinates can be determined. A number of new columns depending on the selected settings (e.g. minimum and maximum distances to the reference set, angle of minimum distance direction etc.) is added to the results table produced by “Measure” on ROI Manager. This analysis was performed on the final StarDist images for PyD and NP-col scaffolds only.

A freehand line on the sample surface was drawn on an adjusted for brightness and contrast Z-projected image of the F-actin channel. XY coordinates of this ROI were saved according to the step-by-step instructions of the macro, and served as the reference file. After obtaining the Results table with the measurements from the ROI manager, the macro was run and on the final StarDist image the lines showing the calculated minimum distances from the nuclei centroids to the sample surface were drawn. The calculated minimum distances were further corrected if needed based on the position of the nuclei regarding the samples surface. For instance, for a calculated distance of 3.67 μm for a nucleus that was above the sample’s surface (the “0” reference), the corrected distance was −3.67 μm.

#### Objects counting – 3D

In this analysis the objective was to obtain measurements for isolated 3D objects of interest, namely the labelled nuclei and actin cytoskeletons, retaining as much as possible their original structure after image segmentation.

##### Nuclei (TO-PRO™-3 staining) – 3D (TrackMate plugin with StarDist detector)

The TO-PRO™-3 channel z-stack was processed for noise reduction, thresholded, and filtered with Gaussian Blur 3D. The TrackMate plugin^3^ was applied on the processed z-stack using the StarDist detector^1^ and LAP tracker for image segmentation following the corresponding instructions.^4^ The obtained segmented Z-stack was added in 3D Manager excluding objects on XYZ edges. The labelled objects were exhaustively verified and treated with Delete, Split, and Merge options as needed to correct discrepancies between the original and the segmented z-stack. In case an object, even after extensive efforts for corrections, was not corresponding to the original z-stack, it was excluded from the analysis. “Measure 3D” was applied to the final objects list.

##### F-actin (phalloidin staining) – 3D (TrackMate plugin with Thresholding detector)

The F-actin channel z-stack was processed for noise reduction. The TrackMate plugin was applied on the processed z-stack defining an appropriate threshold value for the Thresholding detector, and using LAP Tracker for image segmentation. The obtained segmented Z-stack was added in 3D Manager excluding objects on XYZ edges. As for the labelled nuclei, exhaustive verification and corrections took place on the labelled cytoskeletons and all objects not corresponding to the original z-stack, were excluded from the analysis. “Measure 3D” was applied to the final objects list.

#### Proliferating cells position (Ki67 staining) – 2D (StarDist plugin)

The same procedure as described above for the TO-PRO™-3 channel was followed for the Ki67 z-stack. The StarDist plugin was applied on a processed Z-projection of the Ki67 channel and the final segmented image was obtained after the corrections needed. The Results table was also produced as before.

#### Minimum distance: proliferating cells-scaffold’s surface – 2D (Macro Reference Distances)

The same procedure as described above for the TO-PRO™-3 channel was followed for the Ki67 z-stack. The Reference Distances macro was applied on the corrected segmented image from StarDist plugin based on the Ki67 labelling. A final image showing the minimum distances was obtained along with the extended Results table. Again, the calculated minimum distances were further corrected if needed based on the position of the nuclei regarding the samples surface.

#### Colocalization (TO-PRO™-3 – Ki67 staining) – 2D

Before applying Z-projections on the TO-PRO™-3 and Ki67 channels, z-stacks were pre-processed for noise reduction. Z-projections were subjected to Enhance Contrast (saturated pixels 0,3% - Normalize). The final ROI set obtained from the StarDist plugin for the nuclei labelling was applied on the enhanced projections and these areas only were taken into account for the colocalization analysis.

##### Colocalization Colormap

The Colocalization Colormap plugin^5^ was used to obtain a quantitative visualisation of colocalization between processed Z-projections of the TO-PRO™-3 and Ki67 channels.. The Z-projection for the nuclei labelling was always set as “Channel 1” while the Ki67 projection as “Channel 2”. A manual threshold was set on the two images to include all nuclei pixels and the analysis resulted to the corresponding colormap and a correlation index for each examined sample.

### Results

#### Materials for 3D cell culture

**Figure S1.**
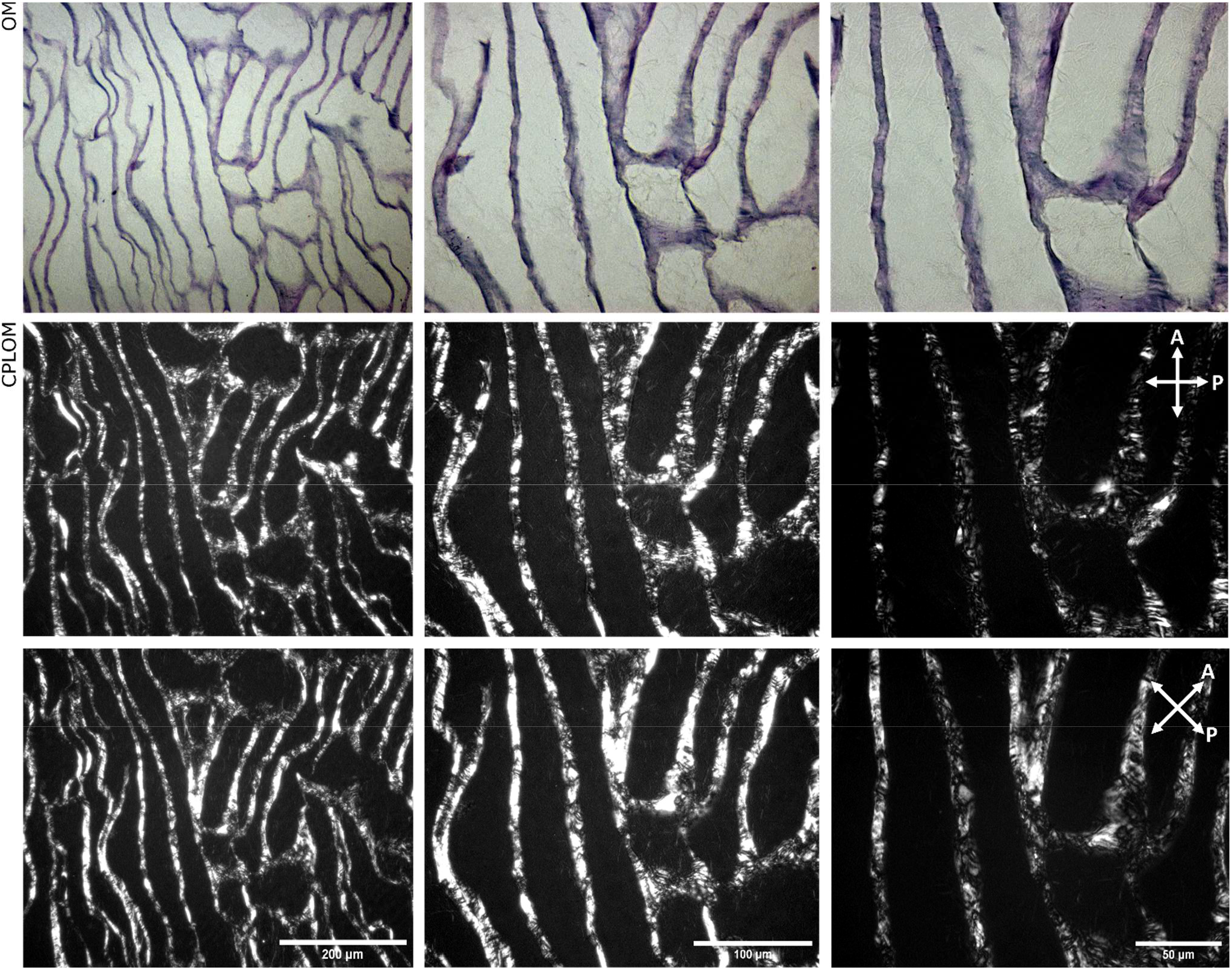
Microscopic characterisation of the PyD collagen matrices at different magnifications. Top row: Optical microscopy images (haematoxylin staining). Middle and bottom rows CPLOM images of the same sections at 0 and 45° between polarizer and horizontal axis, respectively.

**Figure S2.**
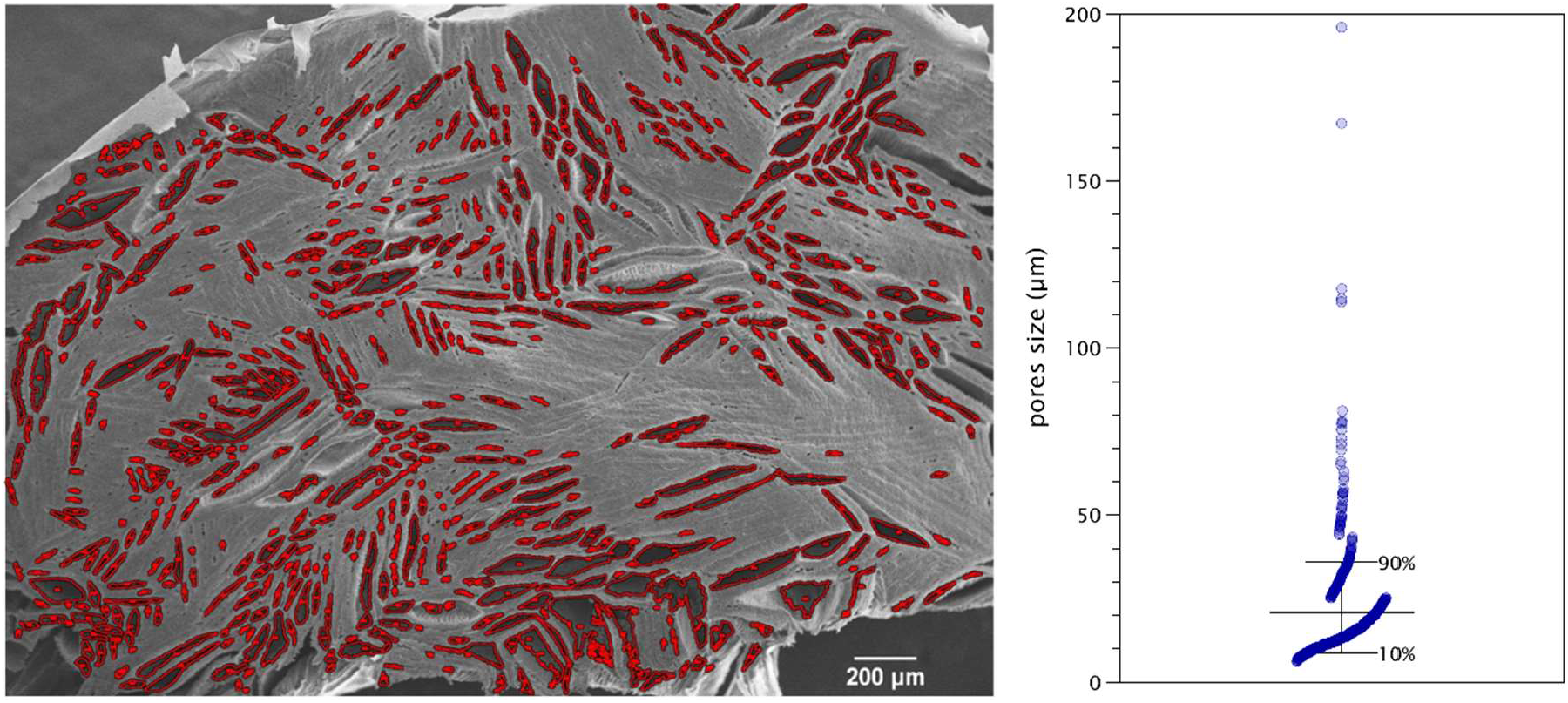
Pore size quantification by Feret minimum diameter.

#### Proliferative status of seeded/encapsulated fibroblasts and colocalization analysis

**Figure S3.**
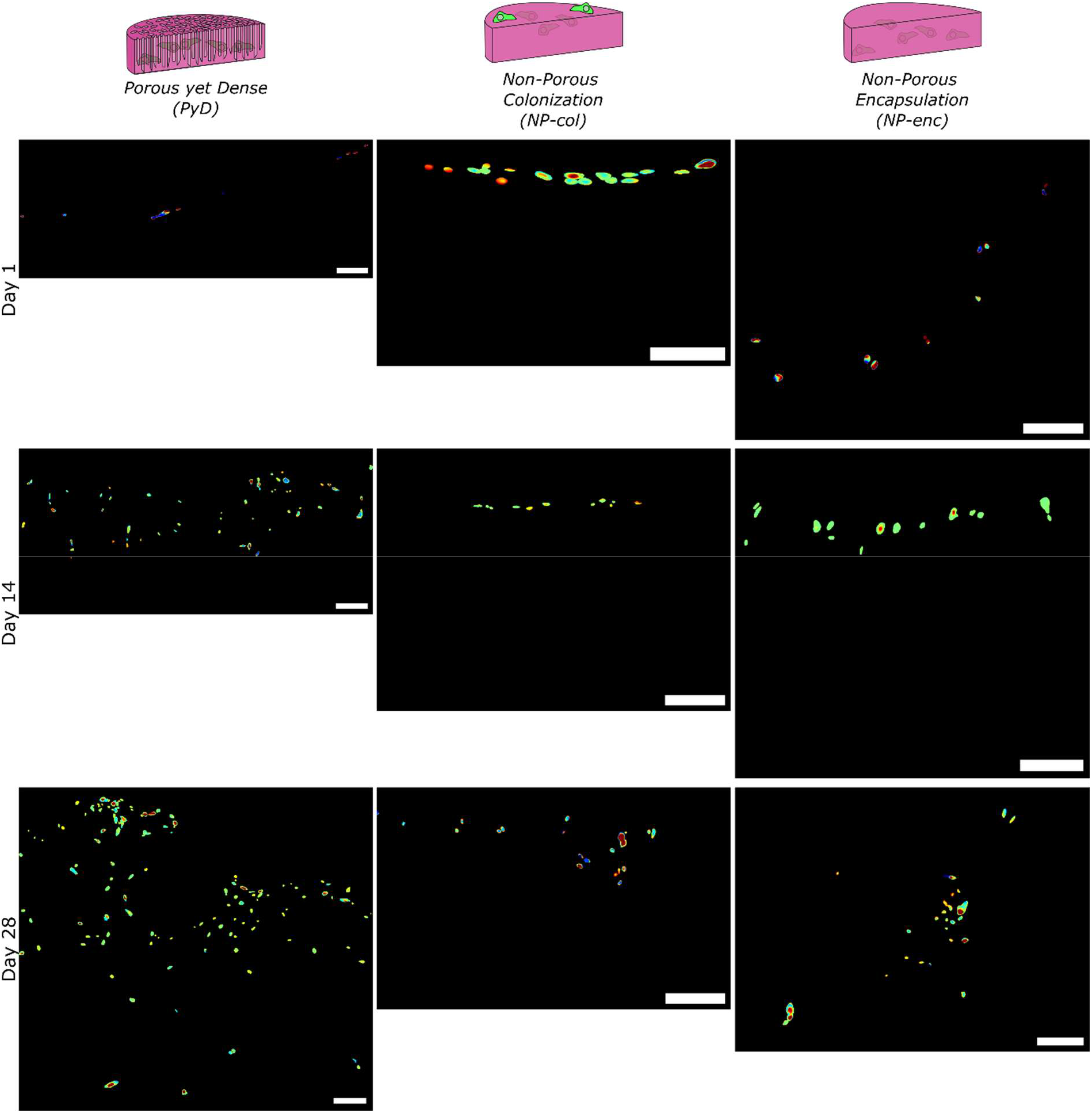
Colocalization colormaps of the processed Z-projections of confocal microscopy images for the TO-PRO™-3 and Ki67 channels. Scale bars: 100 μm.

**Table S1.** Colocalization index values for the colormaps presented in Fig. S3.

|  | PyD | NP-col | NP-enc |
| --- | --- | --- | --- |
| Day 1 | 0.4778 | 0.6790 | 0.6195 |
| Day 14 | 0.5735 | 0.8065 | 0.6987 |
| Day 28 | 0.6919 | 0.5935 | 0.6741 |

#### Morphometric analysis of the seeded/encapsulated fibroblasts

**Figure S4.**
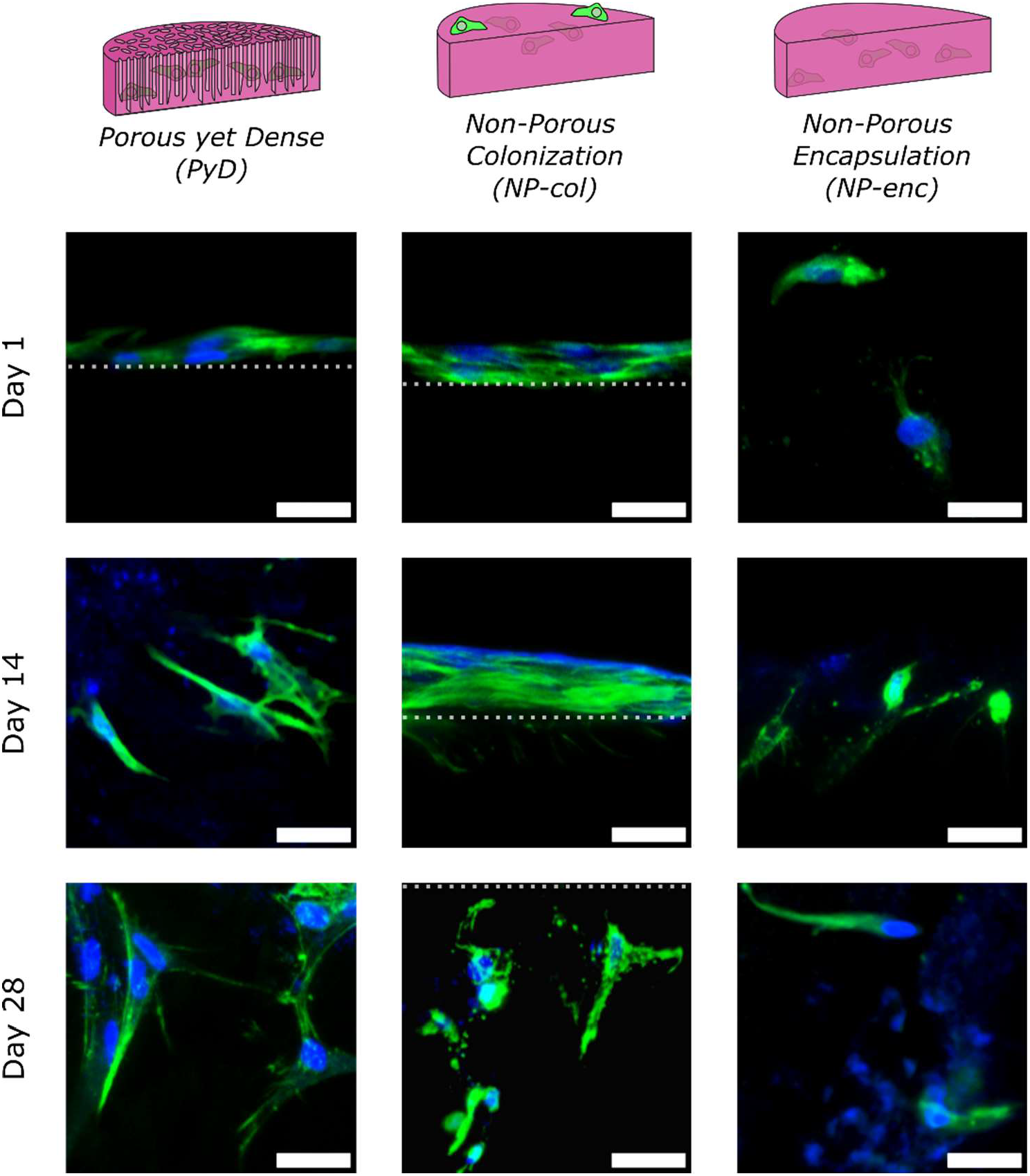
Z-projections of composite confocal microscopy images showing representative examples of NHDFs in the examined samples at the different time points. The dotted lines represent schematically the top surface of the materials. Nuclei: blue (TO-PRO™-3), F-actin: green (phalloidin) Scale bars: 30 μm.

**Figure S5.**
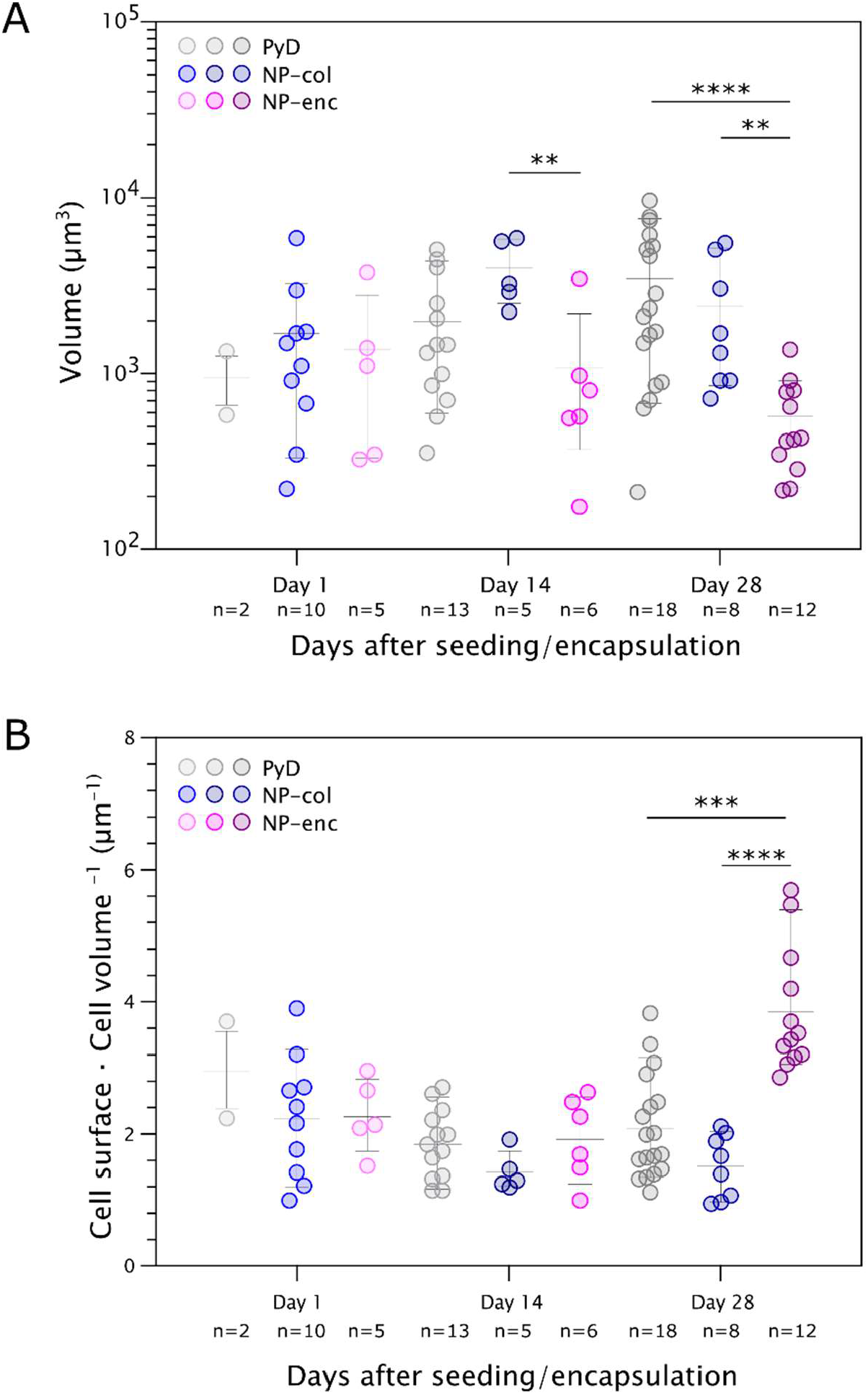
3D morphometric analysis at the cytoskeletal level of fibroblasts colonising PyD and NP collagen materials, based on actin labelling. Scatter plots (mean ± interdecile range plotted) of A) the cellular volume and B) cell surface / cell volume ratio measurements of NHDFs at days 1, 14, and 28 after seeding. Statistically significant differences between different materials for the same time points are shown (**p < 0.0021, ***p < 0.0002, ****p< 0.0001).

**Figure S6.**
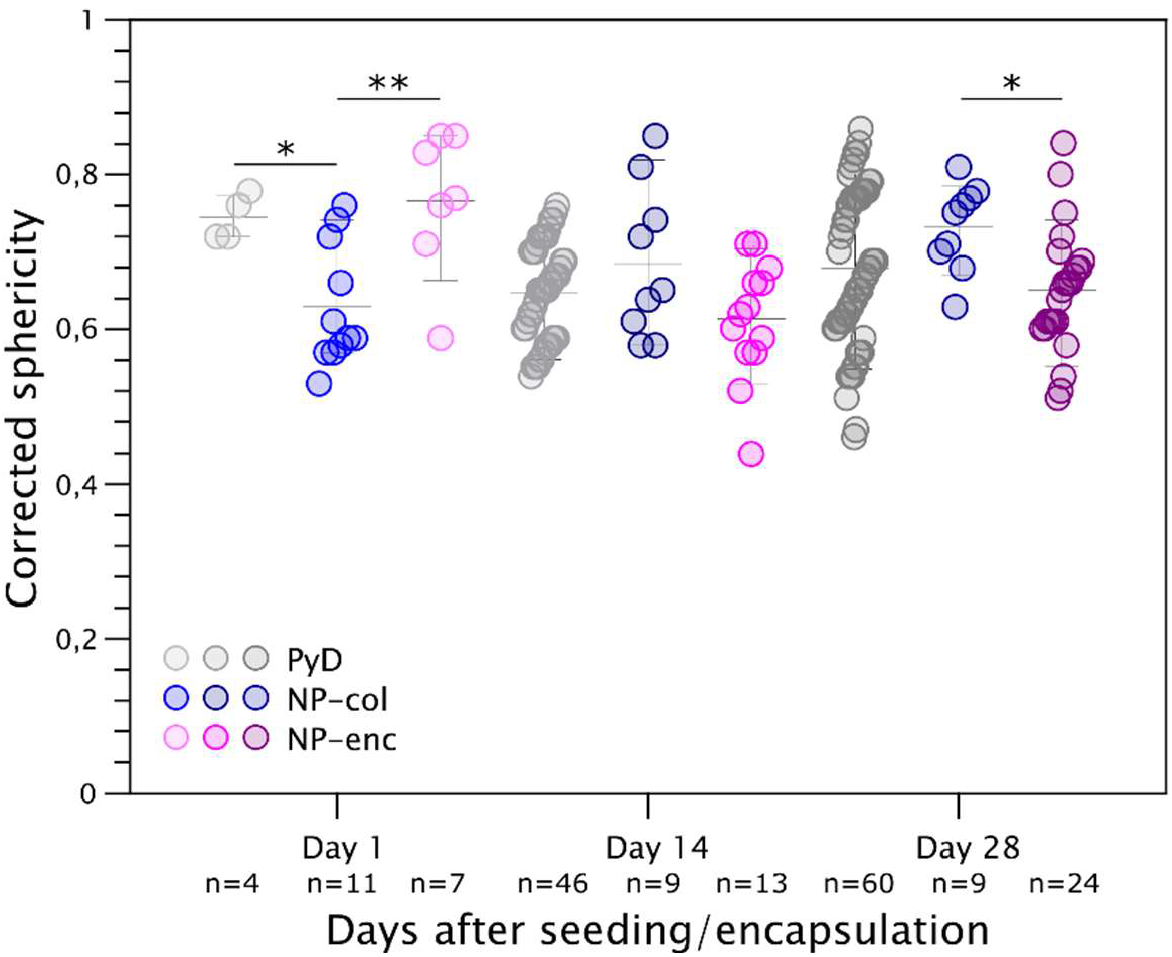
3D morphometric analysis at the nuclear level of fibroblasts colonising PyD and NP collagen materials, based on TO-PRO™-3 labelling. Scatter plot (mean ± interdecile range plotted) of the corrected sphericity measurements of NHDFs nuclei at days 1, 14, and 28 after seeding. Statistically significant differences between different materials for the same time points are shown (*p < 0.0332, **p < 0.0021).

